# Motor protein disruption critically alters organelle trafficking, NMJ formation, and excitation contraction coupling

**DOI:** 10.1101/2025.06.10.658915

**Authors:** H. Bansal, T.A. Hana, A.H. Michael, S. Kamaridinova, J. Bransford, K.G. Ormerod

## Abstract

Trafficking of intracellular cargoes along the neuronal axon microtubule tracks is a motor-protein-dependent process. It is well-established that the motor protein kinesin is responsible for anterograde trafficking of axonal cargo, while the dynein/dynactin complex regulates retrograde trafficking. However, there is still much to uncover regarding the various isoforms of these motor proteins as well as the adapter and cargo-associated proteins involved in the precise trafficking dynamics. Here we use a targeted genetic approach to knockdown candidate kinesin genes involved in trafficking organelles like synaptic vesicles, mitochondria, and dense core vesicles in motor neurons. Using fluorescently tagged cargo proteins; live-imaging experiments were conducted to quantify intracellular trafficking changes, and 2 genes, kinesins 1 and 3, were identified as critical regulators. Disruptions in either gene product, reduce rates of axonal trafficking in motor neurons, and lead to the formation of large intracellular aggregates in somas and axons. Downstream, disruptions in both kinesin 1 and 3 expression led to significant changes in neuropeptide (NP) abundance at boutons, and changes in synaptic morphology, including innervation length, bouton number, and active zone composition. Spinning disc confocal imaging revealed fewer NP trafficking through, or getting captured in kinesin knockdown experiments, and a dramatic reduction in NP release at motor neuron terminals. We go on to show profound reductions in neuromuscular transduction, and excitation-contraction coupling in kinesin 1 knockdowns, but not for kinesin 3. Changes in larval crawling as well as development were observed for kinesin 1 knockdowns. Taken together we have not only identified which kinesins are critically involved in organelle trafficking, but also revealed critical disruptions in cellular morphology, function, physiology, and behavior in genetically disrupted animals.

## Introduction

Motor proteins are critical for transporting intracellular cargoes over long distances using microtubule-assisted tracts along neuronal axons ^1^. Disruption in this transport system can impair neuronal development and function ^2,3^. Neurons uniquely possess axons that can span long distances within the central and peripheral nervous system, which can exceed a meter in length in humans ^4^. While most cellular components are biosynthesized in the soma, it only comprises 1% of the cell volume, while the axon often comprises over 90% of the total cell volume ^5^. Furthermore, intracellular constituents transport bi-directionally, transporting material to synaptic terminals, and subsequently back to the soma, doubling the demand on transport of some cargo ^6,7^. Consequently, the role of trafficking is critical, accentuated by disruptions in axonal transport leading to debilitating diseases such as Alzheimer’s, Charcot-Marie-Tooth, and other neurodegenerative diseases ^8–10^.

Motor proteins are key regulators of long-distance transport in axons ^11^. They require microtubule tracts to move along the axons through an enzymatic process requiring hydrolyzation of ATP in motor-binding domains ^11^. In axons, the plus end of microtubules is oriented towards the periphery of the neuron, near axon terminals. Kinesins travel towards the plus end of the microtubules allowing anterograde transport of cargoes. In contrast, dynein and dynactin mediate the minus-ended transport of cargoes, moving cargo toward the cell soma from axon terminals ^11,12^. Considerable structural and functional variability exists within molecular motors like kinesin, unsurprisingly as more than 45 genes that encode for kinesin superfamily proteins (KIFs) are found in humans, classified into 15 kinesin families ^13–18^. Structurally, kinesins can be broken down into 3 main domains, a motor-domain for microtubule binding via ATP-hydrolysis, a dimerizing stalk domain for structural support, and a cargo-binding domain.

Variability in the stalk and cargo-binding domains enable different kinesin family members to transport different cargo and interact with countless other proteins which can modify their transport speed, accessibility, and mechanochemical properties. Emerging evidence exists to demonstrate that recruitment of different sub-types of kinesins may be organelle or even cargo-specific ^19–24^.

Kinesin proteins are involved in numerous intracellular processes including microtubule dynamics, chromosome alignment, spindle formation, kinetochore assembly, endosome and organelle sorting, and microtubule-based organelle transport^23–27^. Kinesins are known to transport a wide variety of cargo, but most research has focused on trafficking of synaptic vesicles (SVs), dense core vesicles (DCVs), and mitochondria ^19,22–24^. Loss of kinesin-1 disrupts SV transport, causing vesicles to accumulate in the soma rather than reaching synaptic terminals, while mutations in KIF1A, a kinesin-3 family member, result in defective anterograde transport and synaptic dysfunction ^25–27^. Similarly, DCV transport is impaired by KIF1A knockdown, leading to vesicle accumulation in proximal neuronal regions and reduced neuropeptide delivery to axon terminals ^19,28^. In *C. elegans*, mutations in unc^104^, a kinesin-3 homolog, cause DCVs to stall in the soma rather than reach the synapse ^29^. Mitochondrial transport is also dependent on kinesin, as disruptions in Kinesin-1 function lead to mitochondrial clustering in the soma and depletion from axons and dendrites, ultimately affecting energy distribution within neurons ^24,30^. While these studies demonstrate clear defects in intracellular trafficking, the full molecular, cellular, and physiological consequences of these impairments remain incompletely understood.

*Drosophila melanogaster* makes an excellent model for studying motor proteins due to its well-characterized genome and high throughput screening. The fly genome is fully sequenced and highly conserved with many human genes ^31,32^. Additionally, genetic tools such as RNA interference and Gal4/UAS-driven tissue-specific expression allow for manipulation of proteins in vivo ^33,34^. Here we conducted a targeted genetic screen using RNAi to knockdown 6 families of kinesin proteins in *Drosophila* to identify those involved in neuropeptide (NP) trafficking in motor axons. Disruptions in expression in two kinesin families, kinesin 1 and kinesin 3, caused a dramatic effect on NP transport. Live-axonal trafficking experiments reveal both kinesins 1 and 3 are involved in anterograde transport of NPs, while only kinesin 3 disruption impaired retrograde transport. Kinesin 1 knockdown caused numerous effects on neuromuscular junction development, including reducing innervation length, bouton number, and active zone density, while kinesin 3 only impacted active zone density. Kinesin 1 knockdown severely limited the number of DCVs that transported to boutons, and those NPs located at synaptic varicosities had a diminished capacity to move or undergo release from synaptic terminals. Electrophysiological excitatory junctional potentials, muscle force production, and larval crawling were all critically reduced in kinesin 1 knockdown animals, while no significant phenotype was observed for kinesin 3. These findings highlight the important role for kinesin 1 in NP transport and neuromuscular junction development, whereas kinesin 3 primarily contributes to NP trafficking, particularly in retrograde transport, with minimal to no impact on synaptic development and motor behavior.

## Methods

### Fly Stocks and husbandry

*Drosophila melanogaster* flies were cultured on standard media (corn meal, agar, sugar, yeast, and 4-hydroxybenzoate) at 22 °C at constant humidity on a 12:12 light: dark cycle. Fly lines were obtained from Bloomington Drosophila Stock Center (BDSC, see Table 1 for a full list of all fly stocks used).

**Table 1.** Fly lines, genotypes, abbreviations, and BDSC stock numbers used in the current study.

| Genotype | Abbreviation | BDSC Stock # |
| --- | --- | --- |
| OK6-Gal4 | OK6-Gal4 | 64199 |
| UAS-dilp2-GFP | dilp2-GFP | - |
| UAS-khc-RNAi | khc-RNAi | 35770 |
| UAS-Unc<sup>104</sup>-RNAi | Unc-104-RNAi | 58191 |
| UAS-Cana-RNAi | Cana-RNAi | 42640 |
| UAS-Cmet-RNAi | Cmet-RNAi | 35816 |
| UAS-khc-73-RNAi | khc-73-RNAi | 36733 |
| UAS-Kif3c-RNAi | Kif3c-RNAi | 40886 |
| UAS-khc-RNAi | khc-RNAi | 36795 |
| UAS-Klp3A-RNAi | Klp3A-RNAi | 40944 |
| UAS-Pav-RNAi | Pav-RNAi | 42573 |

### Dissection

Wandering stage third-instar larvae were picked from the walls of the vials and dissected at room temperature in calcium-free hemolymph like HL3 saline ^35^. The saline consists of (in mM): 70 NaCl, 5 KCl, 20 MgCl_2_, 10 NaHCO_3_, 115 Sucrose, 5 Trehalose, and 5 HEPES, pH to 7.18-7.19. Dissection was conducted as previously reported ^36^, briefly, the larvae were placed dorsal side up and pinned in the anterior and posterior ends with minutien pins. Using fine scissors, a mid-dorsal cut was made along the length of the animal. The dissected tissue was stretched and pinned down and any viscera was removed using super-fine forceps, leaving the brain and segmental nerve branches intact.

### Fecundity Assay

Motor-neuron selective knock-down of motor-proteins was conducted using RNA interference (RNAi) lines obtained from BDSC (Table 1). UAS-RNAi stock lines for genes of interest were crossed with the motor-neuron specific Gal4-driver, OK6-Gal4. On the day of their eclosion, three males and three virgin females were placed in a vial for three days and then transferred into collection cages placed on-top of 35 mm Petri with grape agar and yeast paste. The grape agar was made with 0.25 mL grape juice, 0.75 mL water, 0.03 g agar, and 0.003 g sucrose. Active dry yeast and DI water were mixed to make the yeast paste. Flies were left in the collecting cages for 26 hours, and afterwards the flies were removed, and the number of eggs were recorded. The eggs were subsequently transferred to a new Petri dish with yeast paste and left for 24 hours. The number of hatched eggs was recorded, and larvae were carefully transferred into a clean food vial. The number of those larvae which matured to third-instar, pupariation, and post eclosion were tabulated.

### Dense Core Vesicle Trafficking

Wandering third instar larvae of each genotype investigated were dissected and filleted, pinned into Sylgard™ bottomed 35 mm Petri dishes in standard HL3.1 saline. Live imaging was conducted using either a Nikon Eclipse microscope, equipped with a Lumencor fluorescence light engine and a Hamamatsu Orca-fusion camera, or a Zeiss Axio Imager 2 microscope with a spinning-disc confocal head (CSU-X1, Yokagawa) and ImagEM X2 EM-CCD camera (Hammamatsu). Live images were acquired using a water-immersion objective, Olympus LUMFL N 60X with a 1.10 NA at either 4 Hz (confocal), or 10 Hz (fluorescence). Videos were exported from either Nikon Elements software or Volocity 3D Image Analysis software (PerkinElmer). Time-series images were imported into Fiji and processed using the Kymograph Clear plugin. DCV time-series recordings were further processed using Kymograph Direct to compute velocity measurements. Velocity was calculated from a minimum of 50 DCVs per animal, and the average velocity of DCVs was plotted for 20 different animals. DCV flux was calculated by drawing a line perpendicular to the axon and quantifying the total number of anterograde and retrograde DCVs moving over the line per unit time. To examine the number of DCVs captured at synaptic terminals, live imaging experiments were conducted on semi-intact animals with the CNS intact. Capture was quantified as DCV entering a synaptic bouton and dropping below the focal plane.

### NMJ and Muscle Immunohistochemical Staining

Immunohistochemistry (IHC) was performed as described previously ^37^. Briefly, dissected third-instar larvae were fixed for 10 minutes or 3 minutes in 4% Paraformaldehyde (PFA) for NMJ and muscle staining, respectively. The larvae were washed 3 times for 10 minutes per wash cycle using 1X Phosphate-Buffered Saline with 0.05% Triton (PBST). The larvae were incubated in primary antibody in PBST at room temperature for 2 hours and washed in PBST three times for 10 minutes. Then larvae were incubated in secondary antibody in PBST at room temperature for 2 hours and washed in PBS three times for 10 minutes. The antibodies employed in this study are as follows: mouse anti-bruchpilot [Developmental Studies Hybridoma Bank (DHSB), stock #nc82, 1:500], phalloidin conjugated with Alexa Fluor^TM^ 488 (ThermoFisher Scientific, catalog #A12379, 1:4000), anti-horseradish peroxidase (anti-HRP, Jackson ImmunoResearch, catalog #123-605-021, 1:500), and goat anti-mouse Alexa Fluor 488 (ThermoFisher Scientific, catalog # A11001, 1:1000). The larvae were mounted on standard microscope slides using glycerol, and a coverslip was placed on top. The larvae were imaged using the Nikon scope described in DCV trafficking above using 60x-water-immersion or 60x-oil-immersion objective lenses. Image analysis was performed using Nikon Elements software.

### Electrophysiology

Larvae were dissected as outlined above in a modified HL3.0 saline with the following composition (in mM): NaCl: 70; KCl: 5; CaCl2:1; MgCl2: 4; NaHCO3: 10; Trehalose: 5; Sucrose: 115; HEPES: 5. (pH was adjusted to 7.18-7.19, daily). Minutien pins were used to pin larvae dorsal side up and the CNS was excised. Motor nerve branches were stimulated (A.M.P.I. Master 8) using a suction electrode at 0.2 Hz. Intracellular voltage of muscle fiber 6 in abdominal segments A2-A4 was recorded using sharp recording electrodes (40–60 MΩ) filled with 3M KCl. Voltage traces were digitized using a 1550 Digitizer (Molecular Devices) and visualized using Axoscope (Molecular Devices) Software. Only recordings with a resting membrane potential of −55 mV to −70 mV were considered for analysis. Excitatory junctional potentials (EJPs) and miniature excitatory postsynaptic potentials (minis) were quantified using ClampFit, and data were exported to Microsoft Excel for data compilation, and subsequently to GraphPad prism for statistical analysis and figure making.

### Force Transducer assay

Force-transducer setup has been described in detail previously ^38^. Briefly, force recordings were obtained using the Aurora Scientific 403A force transducer system (Aurora Scientific, Aurora, Canada), including the force transducer headstage, amplifier and digitizer. A Master 8 (A.M.P.I.) stimulator was used to generated nerve-evoked contractions. The duration of single impulses was 5 × 10−4 s and the interburst duration was kept constant at 15 s. Larvae were dissected as outlined above. To attach larvae to the force transducer, a hook was made from a fine minuten pin and placed onto the posterior end of the larvae. For each animal, 6 replicate contractions were elicited for each animal sequentially, from the following stimulation frequencies:1, 5, 10, 15, 20, 25, 30, 40, 50 100, and 150 Hz. The duration of the burst was kept constant at 600 ms. Digitized data was acquired using Aurora Scientific software, Dynamic Muscle Acquisition Software (DMCv5.5). The digitized data were imported and processed in Matlab using custom code ^38^.

### Larval Crawling Assay

Third instar larvae were isolated from population vials, washed seven times in deionized water, and groups of 5-10 larvae from each genotype were placed in the center of a 1% agar plate for a 5 minute crawling session. Seven recordings per genotype were captured using an infrared camera in a 24″ cubed black box and saved in uncompressed AVI format. The videos were converted to UFMF (Micro Fly Movie Format) using any2ufmf.exe, and CTRAX software extracted CSV files containing X and Y coordinates (in pixels) and heading (in radians). Custom Python and Excel scripts were then used to calculate distance per frame, displacement, instantaneous velocity (using central differences), and linearity (defined as net displacement divided by total displacement over 30-second segments). To account for noise and sudden leaps in position, a Hardle-Steiger sliding-window quantile method was employed for robust smoothing, and outliers were identified both manually and programmatically based on domain-specific criteria. Tracks were segmented by detecting frame gaps and tracking errors, with edge frames trimmed, and mean parameters for each 30-second segment were calculated to reveal genotype-specific locomotor traits.

### Statistical analysis

Analysis was conducted using GraphPad Prism software. For each dataset, the relevant statistical tests were applied, and the results, including F and P values, are presented in the results section. Comparisons were made to control groups unless otherwise specified. Sample sizes were determined based on normality testing. Data are shown as the mean ± SEM (*p < 0.05, **p < 0.01, ***p < 0.001, n.s. = not significant).

## Results

In *Drosophila*, there are at least 10 kinesin families: kinesin 1 (kinesin heavy chain and kinesin light chain), kinesin 2 (Kif3C, Klp64D), kinesin 3 (unc^104^ and KHC-73), kinesin 4 (Klp3A and Kif21A), kinesin 5 (Klp61F), kinesin 6 (pavarotti), kinesin 7 (cmet and cana), kinesin 8 (Klp67A), and kinesin 14 (non-claret disjunctional) ^39^. Kinesins play key roles in several cellular processes such as chromosomal segregation during mitosis, chromosome positioning, microtubule depolymerization, and intracellular trafficking of synaptic vesicles (SVs), neuropeptides (NPs), mitochondria, and organelles ^40–44^. Two families of kinesins, 1 and 3, have been identified and shown to be critical in regulating the trafficking of organelles like SVs, NPs, and mitochondria ^14,18,41,45,46^. We were able to acquire RNAi lines to knock-down 9 different kinesin proteins including 6 different families of kinesins: kinesin 1 (kinesin heavy chain, kinesin light chain), kinesin 2 (Kif3C), kinesin 3 (unc^104^, khc-73), kinesin 4 (Klp3A), kinesin 6 (pav), kinesin 7 (cmet and cana).

### Targeted genetic screen for motors involved in neuropeptide sorting and trafficking

To examine the impact of knocking down kinesin motor proteins on neuropeptide trafficking in motor neurons, fluorescently tagged *Drosophila* insulin like-peptide 2 (UAS-dilp2-GFP; dilp2-GFP) was expressed in motor neurons using the OK6-Gal4 driver ^47,48^. Flies expressing dilp2-GFP, showed a stereotyped expression pattern within the ventral nerve cord (VNC) and segmental nerve branches (SNBs) projecting out to body wall muscles (Figure 1 A). Using a targeted, high-throughput method for screening putative motor proteins involved in neuropeptide sorting and trafficking, OK6-Gal4 > dilp2-GFP was crossed with 9 RNAi lines for candidate kinesin-like peptides (klps) and subsequently, NP aggregation in the VNC and SNBs was examined (Fig. 1). Of the 6 kinesin families, and 9 distinct klps examined, only 3 lines revealed a significant increase in the number of NP aggregation events observed in SNBs. Aggregation was defined as any non-mobile fluorescent puncta, that exceeded 1.25 µm^2^ (Fig. 1 C, One-way ANVOA, F=11.8, P<0.0001). Genetic disruptions in either kinesin 1 genes, khc or klc, resulted in severe breakdowns in NP sorting/intracellular localization within the VNC (Figure 1 A). A quantification of the number of aggregates revealed a ∼5-fold increase in NP-aggregate number compared to controls when knocking down unc^104^, khc, or khc (5.1, 4.8, 4.3, respectively, One-way ANOVA, F=32.4, P<0.0001). The inset in Fig. 1 C shows the density of aggregates for ctl, khc, klc, and unc^104^ (average distance between aggregates in µm, One-way ANOVA, F=10.7, P=0.0001). Next, an assessment of the average size of aggregates was tabulated, and here again, the average size of aggregates for the non-Kinesin 1/ 3 was not significantly different from controls (Fig. 1 D). Knockdown lines for khc, klc and unc^104^ revealed increases in the size of fluorescent aggregates compared to controls, but unc^104^ aggregates were not significantly increased (Fig 1 D, One-way ANOVA, F=11.8, P<0.0001). Given the profound impacts on NP aggregation and localization in the VNC and SNBs, a closer investigation of the dynamics of NP trafficking was necessary for Kinesins 1 and 3. However, of the six kinesin families investigated, only two appear to be critically important in NP localization, and trafficking, kinesin 1 and kinesin 3.

**Figure 1:**
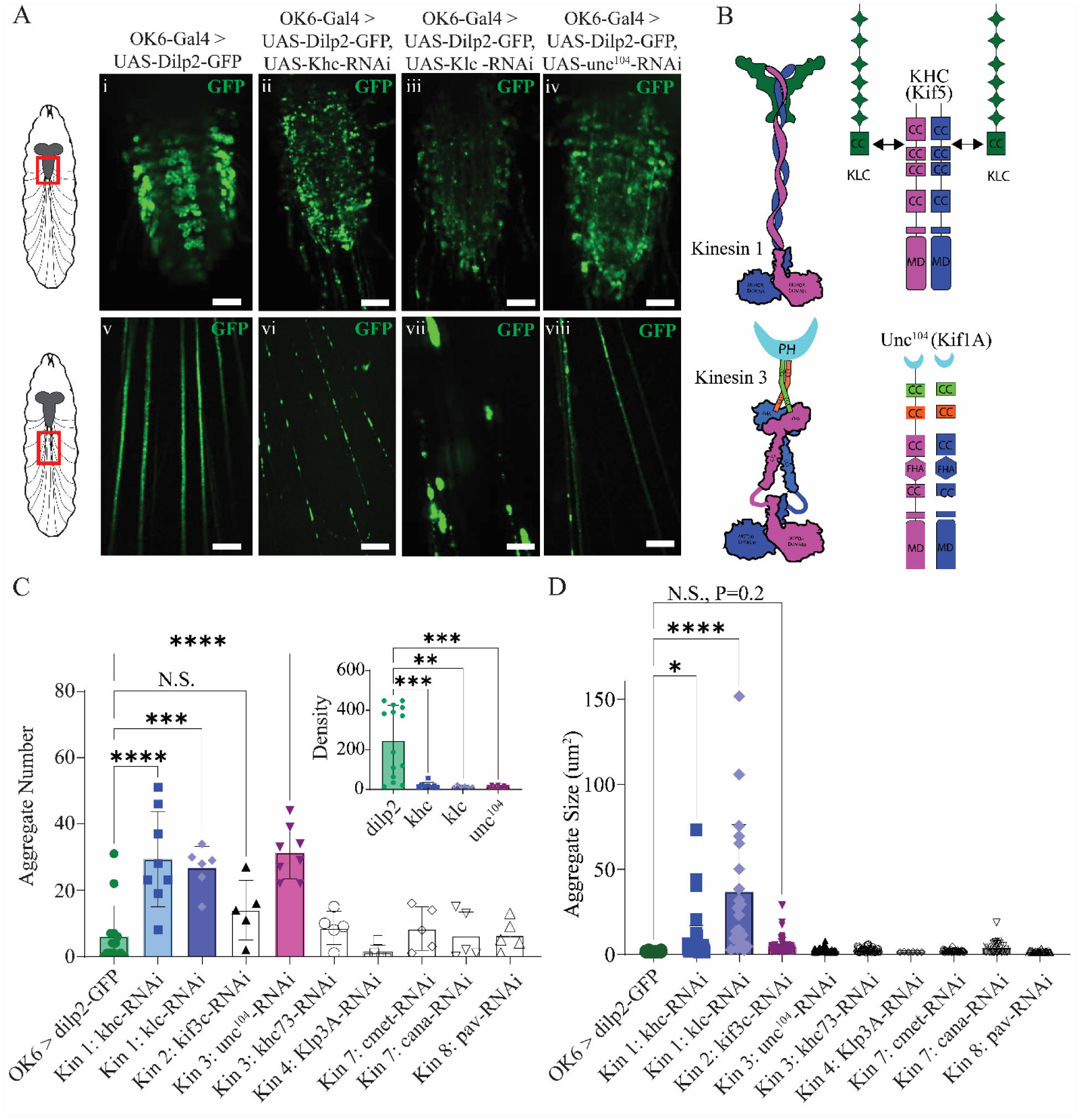
Kinesin disruption critically alters NP trafficking in the nervous system. **A:** Confocal imaging from the ventral nerve cord (VNC), and segmental nerve branches (SNBs), highlighted in cartoon *Drosophila* third-instar larvae using a red box on left inset, top and bottom respectively. **Ai, v:** from control (OK6-Gal4 >dilp2-GFP), **Aii, vi:** kinesin 1 khc knockdown (OK6-Gal4 > dilp2-GFP, khc-RNAi), **Aiii, vii:** kinesin 1 khc knockdown (OK6-Gal4 > dilp2-GFP, klc-RNAi), and **Aiv, viii:** kinesin 3 unc^104^ (OK6-Gal4 > dilp2-GFP, unc^104^–RNAi). Scale bar: 50 µm **B:** Schematic of Kinesin 1 (top) and Kinesin 3 (bottom), highlighting the putative structural organization (left) and functional domains right (cc: coiled coil, md: motor domain, PH: Pleckstrin homology, fha: forkhead-associated). **C:** Quantification of the total number of aggregates from 6 kinesin families and 9 different kinesin related proteins. Inset: density of aggregates in kinesins 1 and 3, units: distance between aggregate in µm. **D:** Average aggregate size for the 9 klps investigated.

### Kinesin knockdown impairs trafficking of neuropeptides

To comprehensively examine the impacts of kinesin 1 and 3 knockdown on molecular trafficking of NPs, live-imaging of NP trafficking in SNBs was conducted using a spinning-disc confocal microscope (Fig. 2). Individual dense core vesicles (DCVs) packaged with fluorescently labelled NPs can be visualized and quantified within axons of SNBs (Fig. 2 A).

**Figure 2.**
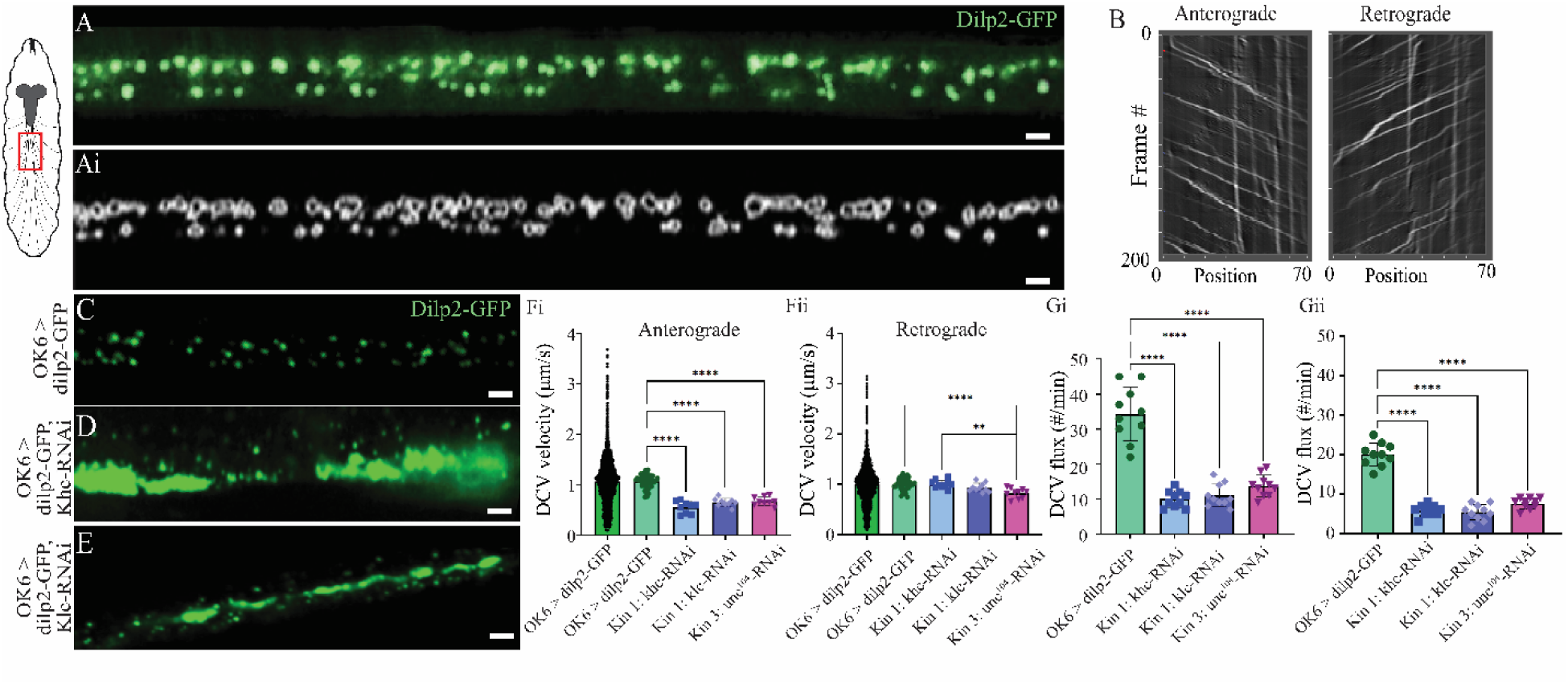
Axonal trafficking of NPs is significantly altered in several kinesin protein knockdowns. *Left:* Schematic of larvae indicating location of imaging. **A**: SNB showing trafficking of the fluorescently tagged NP, Dilp2-GFP. **Ai:** Same axonal tract as A after computational tracking and identification of individual DCVs. Scale bar: 4 µm. **B:** Kymographs created from software Kymograph Direct. Representative SNB images from **C:** OK6 > Dilp2-GFP **D:** OK6 > Dilp2-GFP, khc-RNAi, **E:** OK6 > Dilp2-GFP, khc-RNAi. Scale bar: 4 µm. Quantification of DCV velocity from within one SNB, divided into anterograde (**Fi**) and retrograde (**Fii**). Control data, OK6-dilp2-GFP is demonstrated in two bars. *Left* showing 1000 data points, 50 DCV/animal for 20 animals; *right:* the 50 DCVs per animal averaged for each animal and plotted for the remaining genotypes. **G:** Quantification of DCV flux for the 4 genotypes separated into (**Gi)** anterograde, and (**Gii**) retrograde.

Estimates of DCV velocity were initially measured manually, then subsequently computer software (ilastik) was used to isolate individual DCVs, then track their position from frame-to-frame (Fig. 2 A; Supplementary Videos 1-2). This software exports position data for each frame, for each DCV, from which velocity (um/s), and flux (the number of DCVs moving past a fixed position along the axon per minute) can be extracted. Kymographs were also created using Kymograph Clear and Kymograph Direct (Fig 2 B). Live trafficking of OK6 > dilp2-GFP in SNBs reveal abundant and coordinated antero-and retrograde trafficking (Fig. 2 C, Supplemental Video 1-2). However, live imaging from khc, klc and unc^104^–RNAi knockdown lines, show the severity of NP trafficking, (Fig. 2 D, E). To quantify DCV velocity from OK6 > dilp2-GFP controls, one SNB was recorded from 20 different third-instar larvae. Fifty individual DCVs were measured per SNB, and data were separated into 2 pools, anterograde and retrograde directions, and all 1000 data points were plotted to show the spread of all data (Fig 2 Fi and Fii, left bars). For the remaining bars, data from 50 DCVs were averaged from each SNB from one animal and plotted as a single data point. No significant difference was observed in the rate of DCV trafficking between anterograde and retrograde trafficking in OK6 > dilp2-GFP (Anterograde: 1.07 ± 0.11 µm/s, retrograde: 1.00 ± 0.11 µm/s, T-Test, F=1.2, P=0.2) Knocking-down khc, klc and unc^104^ all significantly reduced the velocity of anterograde DCV trafficking (0.55, 0.65, and 0.68 µm/s, corresponding to 48, 39, and 36% reduction respectively). However, knocking down khc and klc did not significantly impair retrograde trafficking, while surprisingly, knocking down unc^104^ significantly reduce the velocity retrograde DCV trafficking, indicating a potential role for kinesin 3 in retrograde trafficking (Fig. 2 Eii). The total number of DCVs moving both retrograde and anterograde was significantly reduced when knocking down either hc, klc, or unc^104^ (Fig. 2 Gi: Anterograde: One-way ANOVA, F=61.5, P<0.0001, Gii: Anterograde: One-way ANOVA, F=57.1, P<0.0001). While previous investigations have demonstrated a role for these motors in axonal trafficking, there is sparce data exploring the downstream impacts of impaired axonal trafficking on structure, function, physiology, and behavior of the animal.

### Kinesin 1 and 3 motor protein disruption significantly effects growth and development

First, we examined whether knocking down kinesin 1 (khc and klc) or 3 (unc^104^) impacted the growth and development of the animals (Fig. 3). To do so, 3 males and 3 females 72 hours post eclosion were mated for 24 hours, and the total number of eggs laid was tabulated (Fig. 3 A). Significant differences were observed between OK6 - Gal4 control flies, and knocking down khc, klc, and unc^104^, but no significant differences were observed between different kinesin knockdown lines (Fig. 3 C, one-way ANOVA, F=8.7, P<0.001, N=5). After 24 hours the number of eggs that hatched to first instar were tabulated and larvae transferred to fresh food (Fig. 3 A). Knocking down khc significantly reduced the viability of eggs, while khc and unc^104^ knockdowns were not significantly different from controls (Fig 3 D, one-way ANOVA, F=4.9, P=0.01, N=5). The number of first instars the developed to third instars was significantly reduced for both khc and khc knockdown animals compared to control, however no significant impact was observed for unc^104^ animals (Fig. 3 E, one-way ANOVA, F=27.8, P<0.0001, N=5), and similar results were observed for the number of larvae that developed to pupa (Fig. 3 F, one-way ANOVA, F=43.7, P<0.0001, N=5). Tracking the number of eggs that eclosed to adults revealed that knocking down any of the three kinesin transcripts, khc, khc or unc^104^ significantly reduced adults (Fig. 3 G, one-way ANOVA, F=43.0, P<0.0001, N=5). To further assess the health and viability of the animals, the length, width, and area of third instar larvae was examined, however no significant differences we observed for kinesin 1 or 3 knockdown compared to OK6 > dilp2-GFP controls (area: Fig. 3 H, One-way ANOVA; length: F=18.4, width: F=2.9, area: F=7.1, N=10).

**Figure 3.**
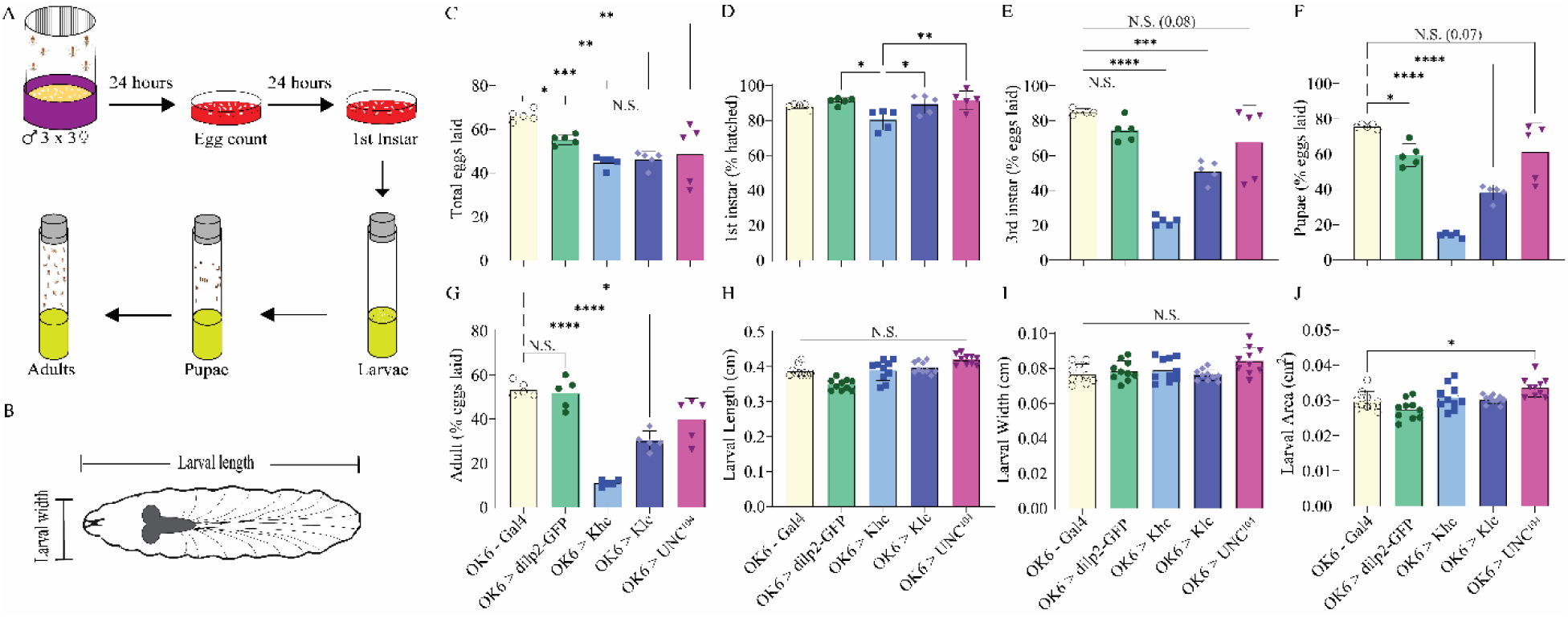
Disruptions in kinesin protein expression significantly alters fecundity but not larval morphology. **A:** Schematic mating and egg-laying assay to track animals throughout development. **B:** Schematic third instar with length and width measurements. For the five genotypes investigated (OK6-Gal4/+: yellow; OK6-Gal4 > UAS-dilp2-GFP: green; OK6-Gal4 > UAS-khc-RNAi: light blue; OK6-Gal4 > UAS-khc-RNAi: dark blue; OK6-Gal4 > UAS-unc^104^– RNAi: pink) the total number of eggs were counted (**C),** the number of eggs that hatched to first-instar (**D),** progressed to third-instar (**E),** pupae (**F),** and adults (**G).** For third-instar larvae, measurements of length (**H),** width (**I),** and area (**J)** are plotted for each genotype.

### Kinesin motor protein knockdown severely impacts motor neuron gross and ultrastructural morphology at the NMJ but not muscles

The larval SNBs projecting from the VNC innervate super-contractile body wall muscles in the periphery responsible for larval movement, like peristaltic locomotion ^38^. There are four different motor neuron subtypes which innervate body wall muscles including the two glutamatergic subtypes Ib and Is, as well as the neuromodulatory type IIs, and lastly the insulin-like immunoreactive type III ^49^. The type Ib and Is (comparable to mammalian tonic and phasic respectively) propagate the output from the CNS underlying rhythmic muscle contractions ^38^. To assess changes in neuromuscular junction (NMJ) morphology and physiology, we first examined the innervation of Ib and Is motor neurons innervating muscle fiber 4 (MF 4, Fig. 4). An assessment of the impact of kinesin knockdowns on MF 4 NMJ was initially conducted by quantifying the number of boutons, size of boutons, and innervation length of the NMJ along the surface of the muscle (Fig. 4, B-E). While there were no significant effects on the NMJ innervation morphology of MN-Is, the NMJ innervation of MN-Ib was significantly reduced for khc knockdown animals (Fig 4 Di, One-way ANOVA, F=8.6, P<0.01). Interestingly, the number of MN-Ib and MN-Is boutons innervating MF 4 was significantly reduced for khc mutants (Fig 4 Ei: One-way ANOVA, F=8.7, P<0.01, Eii: F= 4.2, P<0.05). To change in bouton size was observed for either MN-subtype (Fig. 4 F). To further assess changes in NMJ ultrastructure, immunostaining of bruchpilot (brp), a presynaptic active zone (AZ) scaffolding protein homologous to mammalian ELKS, was conducted ^50^. Both khc and unc^104^ mutants displayed a reduction in brp-positive puncta in MN-Ib boutons (Fig 4, One-way ANOVA, F=6.7, P<0.01).

**Figure 4.**
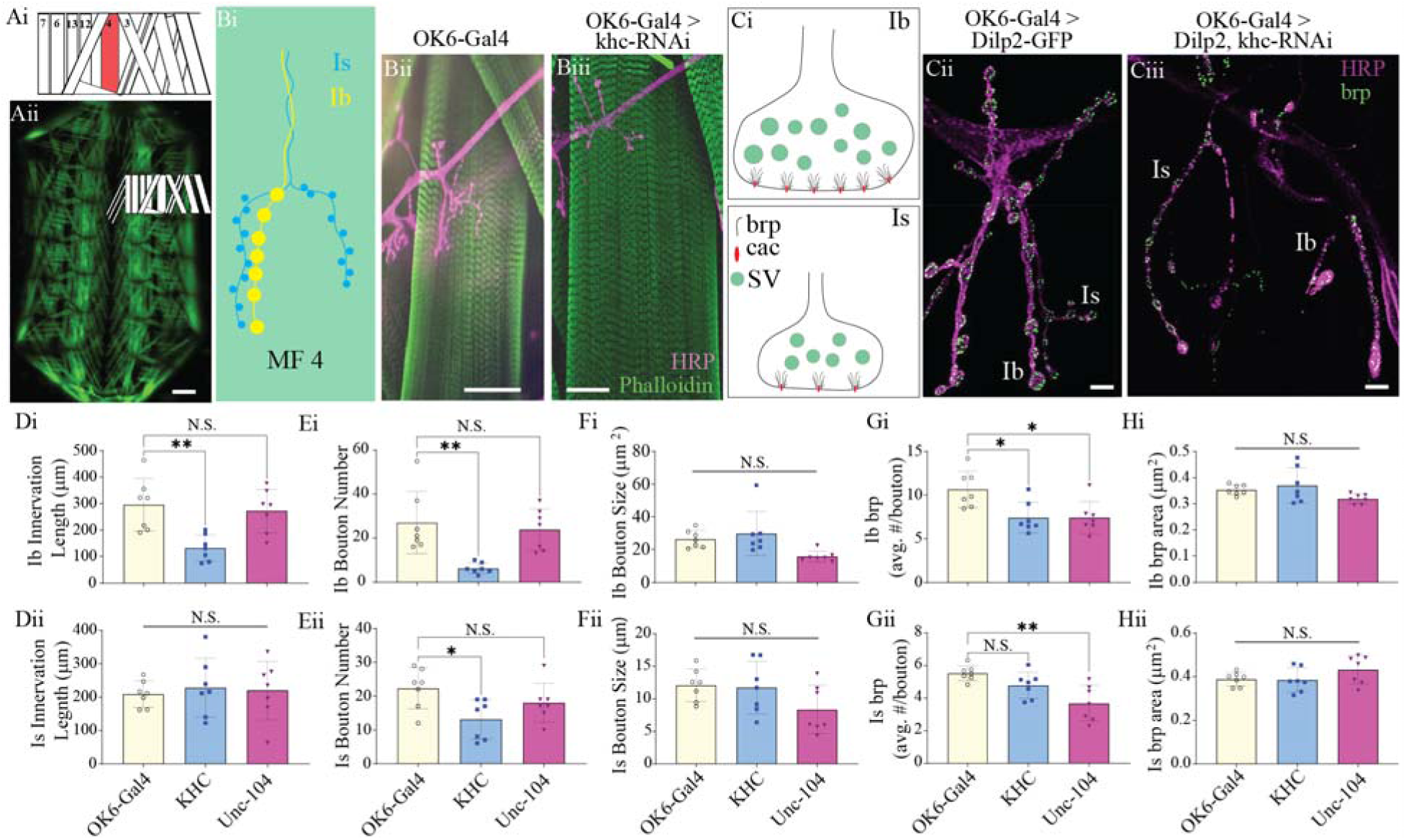
Targeted genetic knockdown of kinesins 1 and 3 significantly impair neuromuscular junction formation. **Ai:** Schematic depiction of third-instar body wall muscles within a single abdominal hemisegment. **Aii:** Fluorescence image of dissected third-instar larvae, highlighting a single abdominal hemisegment in Ai. Scale bar: 500 µm. **Bi:** Schematic depiction of glutamatergic Ib and Is innervation along the surface of muscle fiber 4. **Bii:** Immunostain from OK6-Gal4 and **Biii:** OK6-Gal4 > UAS-khc-RNAi: with HRP: purple, and phalloidin: green. Scale bar: 50 µm. **Ci:** Schematic glutamatergic Ib (top) and Is (bottom) boutons depicting bruchpilot (brp), cacophony (cac), and synaptic vesicles (SV). **Cii:** immunostain from OK6-Gal4 > UAS-dilp2-GFP: green; and **Ciii:** OK6-Gal4 > UAS-khc-RNAi: with HRP: purple, and DCV: green. Scale bar: 4 µm. Quantification of changes in innervation length **(D),** bouton number **(E),** bouton size (**F)**, brp density (average number per bouton, **G)**, and average BRP area (**H)**, separated by Ib (i, top) and Is (ii, bottom) MN subtypes.

However, only unc^104^ knockdown animals displayed a reduction in brp-positive puncta in MN-Is boutons, while khc-knockdown animals bordered on significance with P=0.056 (Fig 4, One-way ANOVA, F=8.8, P<0.01). Given the substantive changes in motor neuron morphology and the intimate and dynamic relationship occurring between motor neurons and body wall muscles during embryonic development, we next assessed whether knocking down motor proteins in motor axons impacted muscle ultrastructure (Fig. 5). First, an assessment in gross muscle morphology was conducted by examining changes in individual muscle fiber length, width, and area, but no significant differences were observed (Fig 5, One-way ANOVAs: length: F=0.7, P=0.5; width: F=0.4, P=0.7; area: F= 0.7, P= 0.5). Next, a fluorescence line-profile was conducted using phalloidin-stained muscles to approximate the length of sarcomeres and I-bands, but no significant differences in muscle ultrastructure was observed (Fig. 5, One-way ANOVA, sarcomere: F=0.4, P=0.7, N=8, I-band: F, P=0.7, N=5).

**Figure 5.**
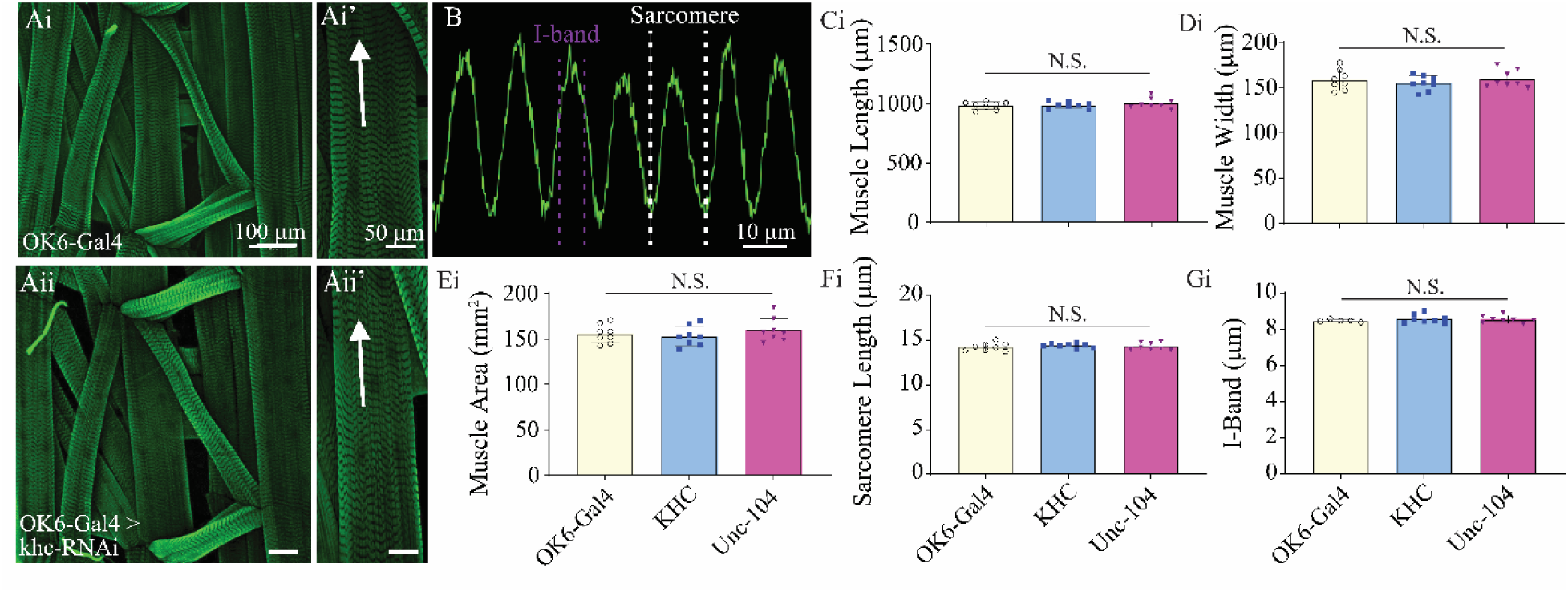
Muscle ultrastructure is not altered in kinesin 1 or 3 knockdown lines. Immunostains from dissected larvae showing body wall muscles in a single abdominal hemisegment from (**Ai**) OK6-Gal4, and (**Aii**) OK6-Gal4 > UAS-khc-RNAi. **Ai’** and **Aii’** depict confocal images highlighting MF 4 from OK6-Gal4 and OK6-Gal4 > UAS-khc-RNAi, respectively. Arrow indicates location and direction of fluorescence intensity profile line. **B:** Fluorescence intensity profile line for GFP (phalloidin). Dashed white and magenta lines indicate how sarcomere and I-band measurements were calculated respectively. Quantification of muscle length (**C**), muscle width (**D), muscle** area (**E),** sarcomere length (**F**), and I-band (**G**).

### Knocking down kinesins 1 and 3 dramatically reduces NP abundance, movement, and release at the NMJ

During the initial screening of motor protein knockdown on NP aggregation and localization (Fig. 1), a substantial change in fluorescence localization in the periphery and at NMJs was noted. Representative confocal images from OK6-Gal4 > dilp2-GFP show a stereotypical distribution of DCVs both within axons and at motor neuron terminals (Fig 6 Ai-ii). However, knocking down khc results in a dramatic reduction in NP abundance in motor neurons along the surface of muscles and at synaptic varicosities (Fig. 6 B). A co-stain for axonal membrane was conducted with anti-HRP to quantify average GFP fluorescence within axons and at NMJs along the surface of MF 4, and a significant reduction was observed for both khc and unc^104^ knockdown animals (Fig 6 D: OKD-Gal4: 920 ± 351, OK6-Gal4> dilp2-GFP, khc-RNAi: 228 ± 63, OK6-Gal4> dilp2-GFP, unc^104^–RNAi: 314 ± 101, One-way ANOVA, F=15.4, P<0.0001). Using spinning disc confocal microscopy, the movement of DCVs entering a non-terminal bouton was assessed. For OK6-Gal4 > dilp2-GFP animals, an average of 3.6 ± 1.4 DCVs per minute while for khc and unc^104^ animals an average of 0.4 ± 0.5 and 0.3 ± 0.5 per minute, respectfully were observed (Fig. 6 E, One-way ANOVA, F=45.5, P<0.0001, N=10). Next, to assess rates of synaptic capture 50 DCVs were observed per bouton, and the total number that transited through vs were captured into boutons was counted and converted to a percentage. For OK6-Gal4 > dilp2-GFP animals, 83.2 ± 2.1 % of DCVs transited through the boutons. Given that so few DCVs were mobile, and their velocity of trafficking was severely reduced, very few DCVs transited through boutons during the experiments. Consequently, for the 3-4 animals where DCV mobilization was observed through a bouton, none were captured resulting in significant differences compared to controls for both khc and unc^104^ knockdown animals (One-way ANOVA, F=3417, P<0.0001, N=3-10). Lastly, to determine if knocking down kinesin 1 or 3 impaired DCV release from boutons a ‘fusion-assay’ was conducted using previously published approaches ^51^. Axons from motor neurons were stimulated at 70 Hz stimulation for 20 seconds, and DCV exocytosis was quantified as a reduction in GFP fluorescence at Ib boutons (Fig 6. Ci-ii). However, when directly stimulating boutons from either kinesin 1 or kinesin 3 knockdown animals, a significantly impaired reduction in stimulation-induced fluorescence change was observed (Fig 6 G). Any drop in fluorescence is likely a reflection of photobleaching given that no redistribution in DCV was observed, as was observed for controls (Fig 6. Ei vs Eii).

**Figure 6.**
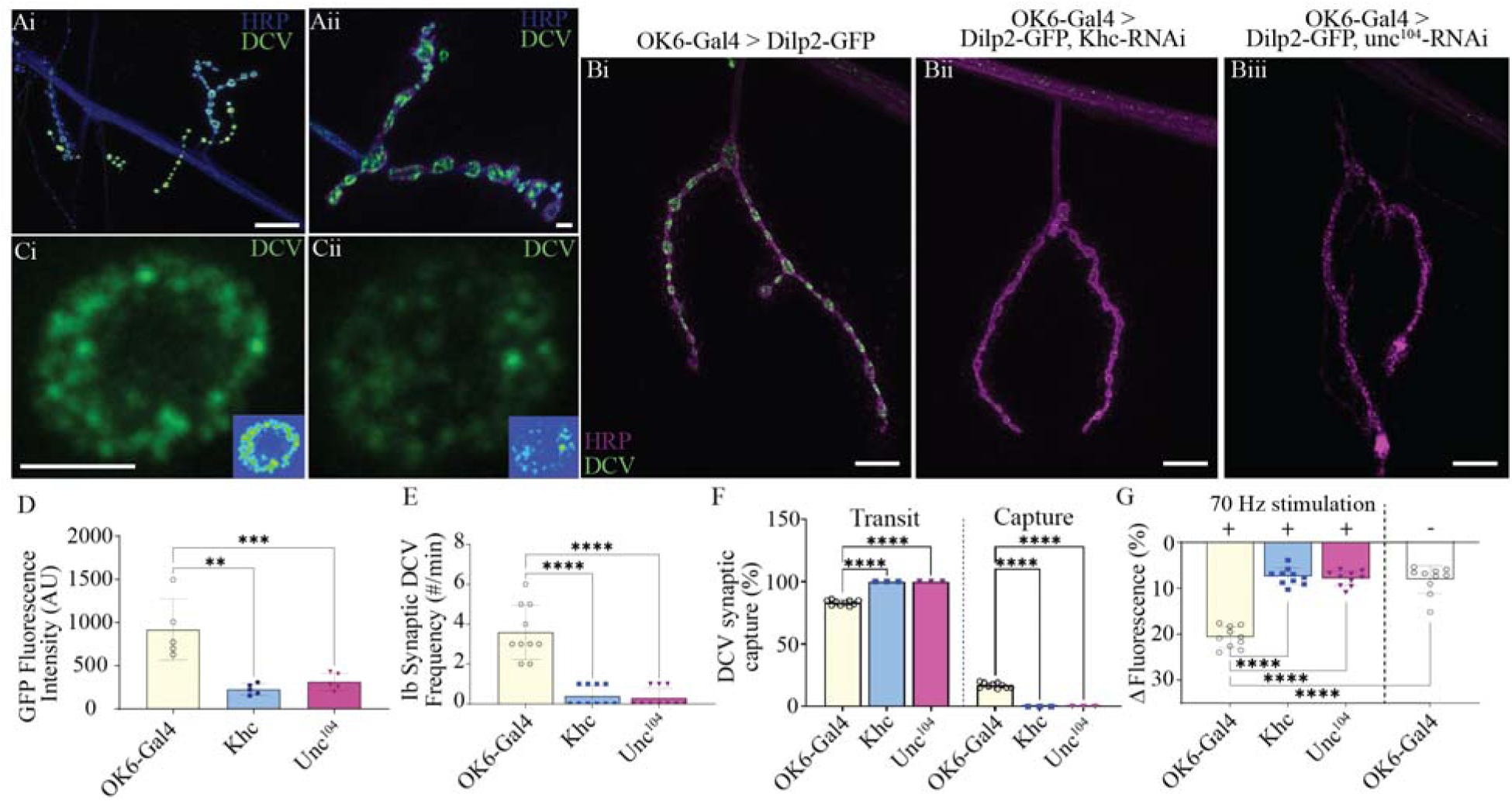
Motor protein knockdown profoundly impacts DCV mobility and release at synaptic boutons. Immunostain images showing (**Ai**) peripheral nerve branches and multiple strings of synaptic varicosities, and (**Ai’**) zoomed-in image of a single string of synaptic varicosities from OK6-Gal4 > dilp2-GFP, stained with HRP: blue, and DCVs: green. A representative image from a single bouton taken during live spinning-disc confocal imaging before (**Bi**) and after (**Bii**) high frequency, 70 Hz stimulation. *Inset*s: heatmap showing fluorescence intensity. Immunostains to show changes in DCV localization at synaptic varicosities in (**Ci**) OK6-Gal4 > UAS-dilp2-GFP: (**Ci**) OK6-Gal4 > dilp2-GFP, khc-RNAi, (**Ci**) OK6-Gal4 > dilp2-GFP, unc^104^–RNAi. Quantification of: (**D**) changes in GFP fluorescence at synaptic varicosities, (**E**) frequency of DCVs at boutons, (**F**) number of DCVs transiting through ‘transiting’ and those entering bouton “synaptic capture” (**G**) drop of GFP fluorescence at non-terminal boutons following 70 Hz stimulation for 20 s.

### Kinesin 1, but not 3, alters neuromuscular transduction

Given the dramatic changes in NMJ ultrastructure and NP mobility and release, we next assessed whether knocking down kinesin 1 or kinesin 3 impacted neuromuscular transmission (Fig. 7). These molecular motors are known to transport not only NPs via DCVs, but other organelles like synaptic vesicles (SVs), and mitochondria. Using sharp intracellular recordings, a dramatic reduction in excitatory junctional potentials (EJPs) was observed for kinesin 1 knockdown animals, but not for kinesin 3 knockdowns compared to controls (Fig. 7. A, C, One-way ANOVA, F=1812, P<0.0001, N=7). No observable change was found for miniature excitatory junctional potentials (Fig. 6 B, minis) amplitude (Fig. 7 D, One-way ANOVA, F=0.04, P=0.9, N=7), however significant reduction was observed for both kinesin 1 and 3 knockdowns (Fig. 7 E, One-way ANOVA, F=6.7, P=0.01, N=7).

**Figure 7:**
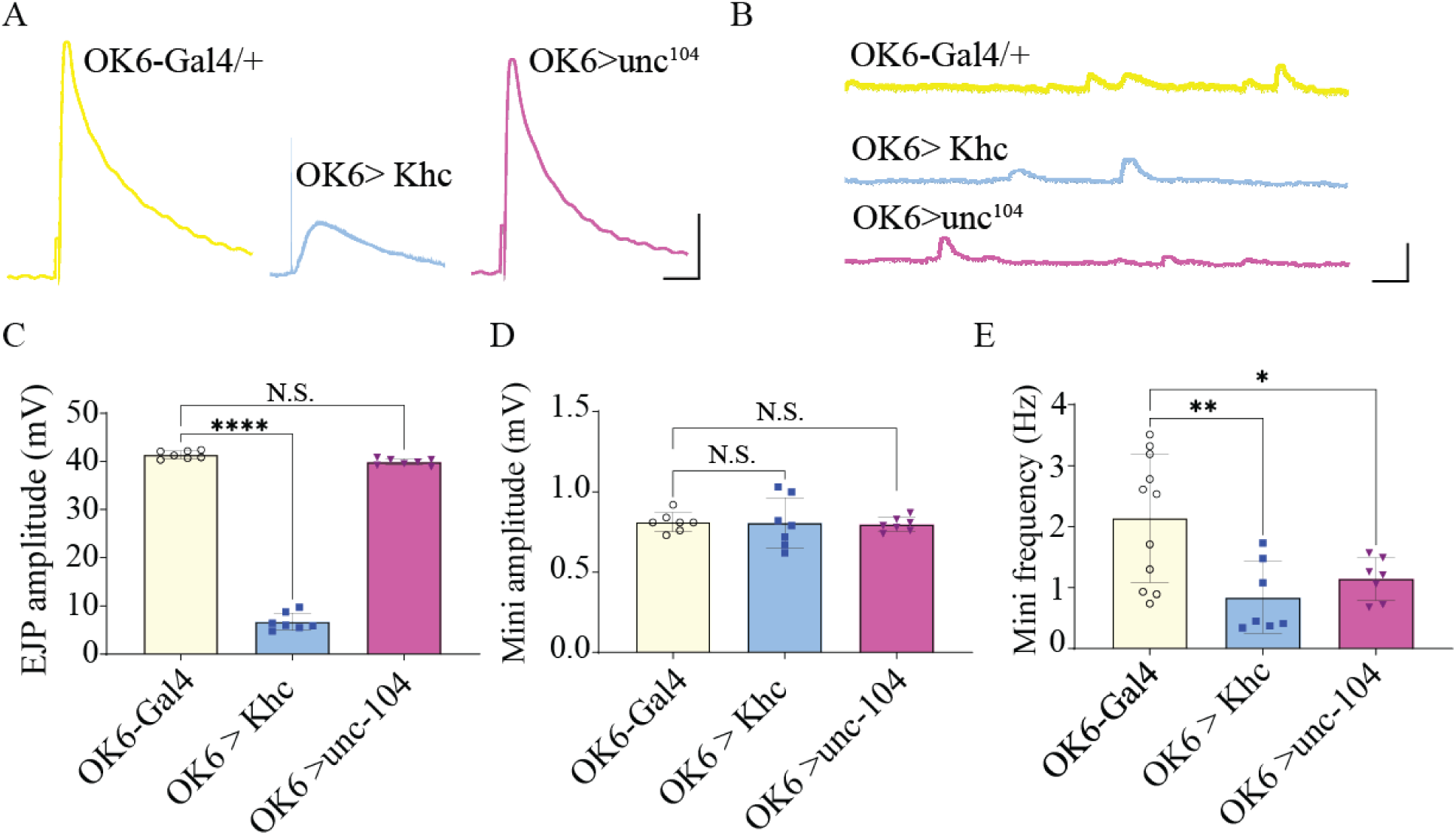
Neuromuscular transduction is significantly impaired only in kinesin 1 knockdown lines. Representative electrophysiological recordings from the three genotypes showing excitatory junctional potentials (**A,** EJPs) and miniature end plate potentials (**B**, minis). Quantification of changes in (**C**) EJP amplitude, (**D**) mini amplitude (**E**) mini frequency.

### Kinesins 1, but not 3, reduce excitation-contraction coupling

To further examine how kinesin 1 and 3 knockdown in motor neurons would impact the NMJ, an examination of muscle force production from larval body wall muscle was explored. We previously established methodology to examine muscle force production by generation a force-frequency curve, spanning the entirety of the endogenous motor neuron frequencies, ranging from 1-150 Hz ^38^ (Fig 8. A-D). Knocking down kinesin 3 did not impair muscle force production at any point along the force-frequency curve, nor was maximal force production reduced compared to controls (Fig. 8 E-F). However, knocking down kinesin 1 dramatically reduced muscle force production at every stimulation frequency investigated (Fig. 8 E). Unlike control and kinesin 3 knockdown animals which produced muscle force from a single AP, kinesin 1 knockdown animals did not generate quantifiable force until 20 Hz stimulation was delivered (Fig. 8 E). The reduction in force generation was greater than 80% above 50 Hz representing one of the most dramatic phenotypes observed using this experimental approach^37,38,52^. The maximum force generated by kinesin 3 knockdown animals was not significantly different from controls, however, a significant reduction was observed following knockdown of kinesin 1 (Fig. 8 F, One-way ANOVA, F=64.2, P<0.0001, N=8).

**Figure 8:**
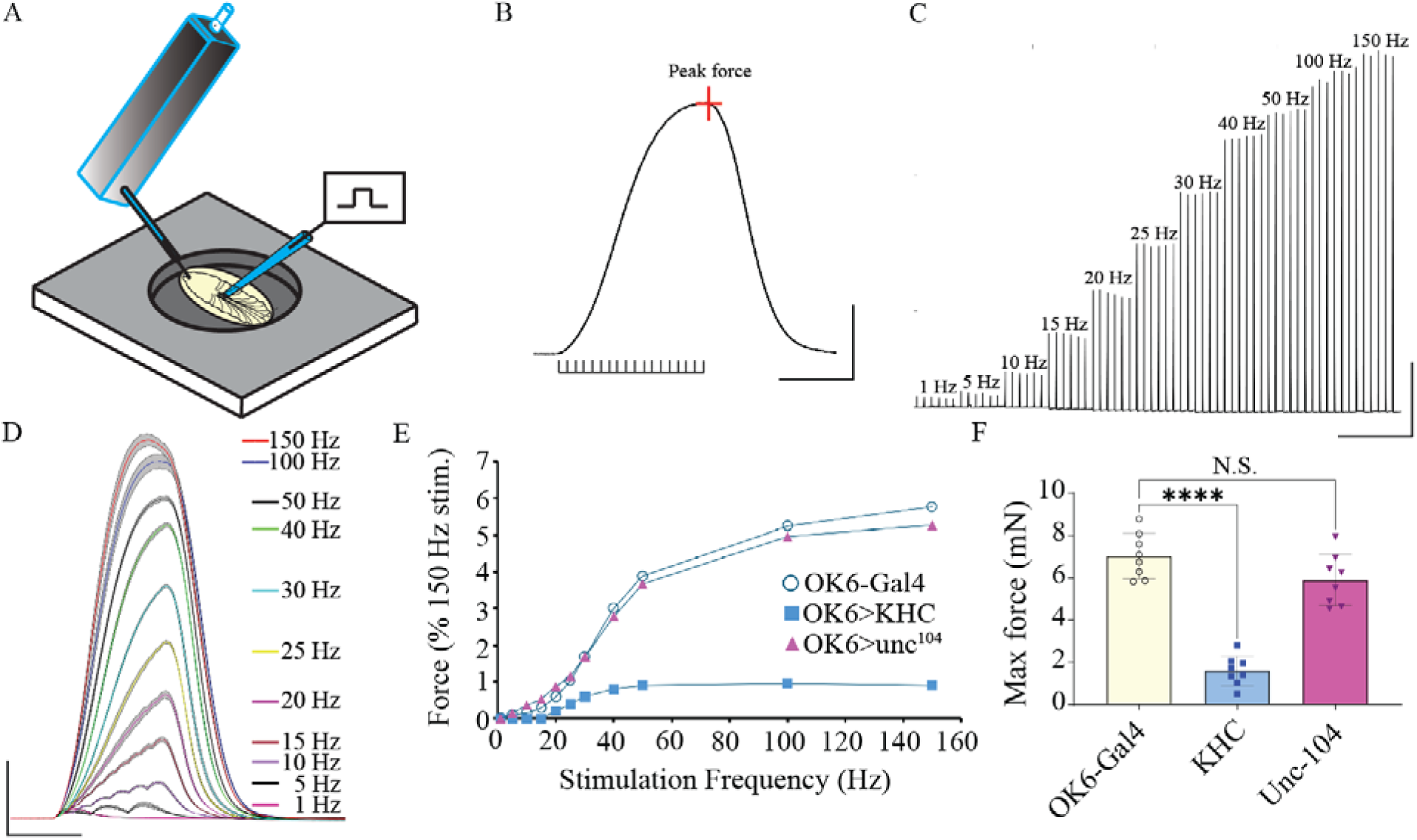
Excitation contraction coupling is severely reduced in kinesin 1 knockdown larvae. **A:** Schematic representation of muscle force transducer setup. **B:** A single muscle contraction trace induced by a 40 Hz stimulation for 600 ms. **C:** larval motoneuron stimulation protocol used to generate dynamic force-frequency recordings for muscle contraction from initial threshold through saturation (1–150 Hz). Stimulus duration was kept constant at 600 ms. D: The six replicate contractions from each stimulation frequency shown in **C** were averaged and plotted with 95% confidence interval (grey area). CI. **E:** Force-frequency plot from each of the 3 genotypes. **F:** Quantification of maximal force generated from 150 Hz stimulation for each of th 3 genotypes.

### Disruptions in kinesin 1, but not 3, result in changes in larval behavior

Lastly, to assess whole-organism behavioral changes resulting from kinesins 1 and 3 knockdowns, larval crawling was conducted. Underlying larval crawling are central pattern generators sending high frequency patterned output from the ventral nerve cord to the muscles from the CNS via motor neurons in the VNC 55. Consistent with electrophysiological and force recordings, kinesin 1 knockdown significantly impacted larval crawling across numerous metrics examined (Fig. 9). Ctrax was used to track larvae, and custom code was created to track quantify crawling, enabling the generation of crawling patterns (Fig. 9 Ai-iii). From these, linear velocity and linear displacement (Fig 9 B, Fig. 9 C, Brown-Fosythe ANOVA, F=9.7, P<0.01), and linearity index (Fig. 9 D, Brown-Fosythe ANOVA, F=10.3, P<0.01) were all significantly reduced from kinesin 1 knockdown animals, but not kinesin 3 (Suppl. Vid 3-5).

**Figure 9:**
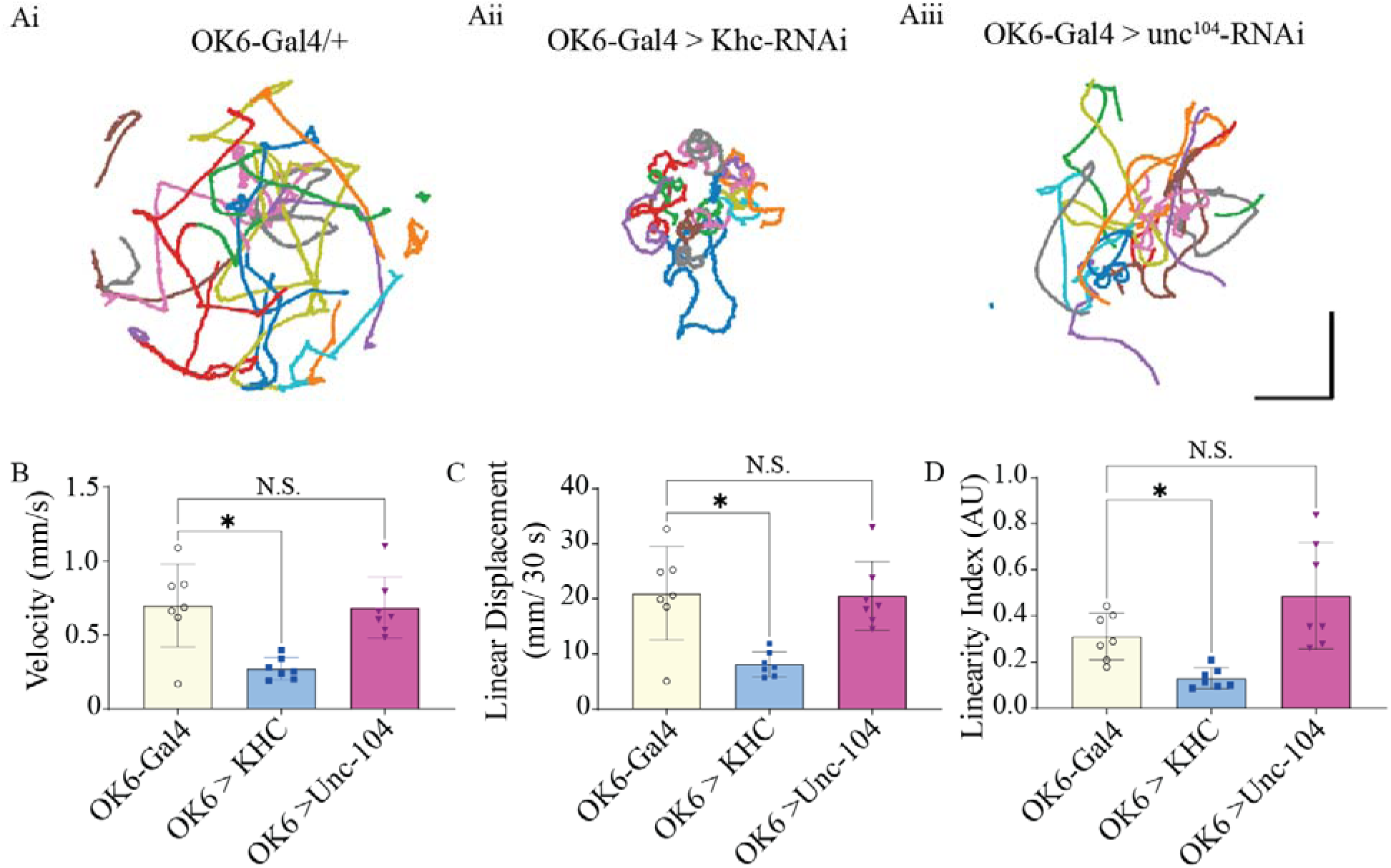
Deficits in larval crawling behavior are only observed in kinesin 1 knockdown animals. Schematic representing larval crawling behavioral changes from (**Ai**) OK6-Gal4/+; (**Ai**) OK6-Gal4 > khc-RNAi, and (**Ai**) OK6-Gal4 > unc^104^–RNAi. B, C, D: Seven videos of 10 larvae from each genotype were quantified and the positional data were averaged and plotted at a confidence interval of 95%; velocity (B) was measured as millimeters per second. Linear displacement (C) was calculated as *d* = √((x_end_ - x_start_)² + (y_end_ - y_start_)²) per 30 second segment of larval track, then averaged. Linearity index (D) was measured as the linear displacement divided by the total displacement (nonlinear), providing an index of total time spent turning vs. actual distance traveled.

## Discussion

Here we conducted a targeted screen of six kinesin families, encompassing nine klps, to assess their role in the trafficking of neuropeptides and the downstream impacts of kinesin disruption in motor axons. Upon screening nine candidate kinesins involved in neuropeptide transport, we identified two kinesins, 1 and 3, that produced an aggregation phenotype. This finding aligns with prior studies demonstrating the essential role these kinesins in transport of other NPs, islet cell diabetes autoantigen-1 in *C. elegans* and atrial natriuretic factor (ANF) in *Drosophila* ^53,54^. Other kinesins, including Kif3C, KHC-73, KLP3A, CENP-meta (cmet), CENP-ana (cana), and Pavarotti, did not produce an obvious aggregation phenotype in the VNC or SNBs. This outcome was unsurprising, as Kif3C has been demonstrated as a microtubule-destabilizing factor required for axon regeneration rather than cargo transport ^55^. Cana and cmet, *Drosophila* homolog of CENP-E, play key roles during stages of cell division, mediating processes such as chromosome alignment, spindle formation, and kinetochore assembly ^43,56,57^. In mice, KLP3A mutations are understood to disrupt central spindle and cytokinesis in males, although somatic cells in *Drosophila* show insensitivity to loss of wild-type KLP3A ^58^. On the other hand, KHC-73 has been shown to facilitate the routing of synaptic endosomes back to the soma ^59^. Finally, Pavarotti forms the centralspindlin complex, which plays a role during cytokinesis ^60^. Both kinesins, 1 and 3 exhibited a significant increase in the number of neuropeptide (NP) aggregates within SNBs, with kinesin 1 also showing a significant increase in aggregate size. While there are at least 10 families of kinesins in *Drosophila* to date our data supports previous observations of kinesin 1 and 3 as pivotal motors of axonal NP trafficking ^11,12,61–63^.

Given that knocking down kinesin 1 and 3 led to significant accumulation of NP in somas and axons, we next explored their direct role in trafficking. Our investigation revealed that while both kinesins 1 and 3 are required for efficient anterograde transport, only kinesin 3, unc^104^, plays a role in retrograde transport. Previously, retrograde transport of ANF filled DCVs was significantly disrupted in unc^104^ mutants with no observable defects in retrograde transport of synaptic vesicles and mitochondria ^19^. One possibility for this kinesin-dependent role in retrograde transport lies in the complex of proteins associated with different organelles. Mitochondria have a unique adapter protein Milton or Observations of Kif1C in mice, a kinesin 3 member, has shown its association with adaptor protein, HOOK3, whose binding enables dynein-dynactin recruitment, thereby facilitating retrograde transport ^64^. Additionally, the phosphorylation of other adapter proteins huntingtin and JNK-interacting proteins (JIP) are known to mediate the switch between retrograde and anterograde transport of cargo, which differentially interact with kinesin subtypes ^65,66^.

Intracellular organelle and protein aggregation in the nucleus and axons of neurons disrupts neuronal homeostasis, leading to cytotoxicity, neurodegeneration, and ultimately neuronal necrosis/apoptosis. Much of our current understanding of the downstream implications of protein aggregates is predicated based on neurodegenerative diseases like Amyotrophic lateral sclerosis (ALS), Alzheimer’s (AD), Parkinson’s (PD), and Huntington’s disease (HD) ^6,7,52^. Consequently, we explored the downstream ramifications of altered kinesin motor protein activity selectively in motor neurons. Initially, we investigated the effects of kinesin 1 (khc) and 3 (unc^104^) knockdown on the growth and development of *Drosophila*. A morphological assessment of third-instar larvae did not reveal any significant changes in larval length, width, or area. This was surprising given previous reports from khc and klc mutant *Drosophila* larvae describe severe morphologically and physiologically impaired second and third instars ^67–70^. By tracking multiple developmental stages, from egg laying to adult eclosion, we aimed to determine whether impaired kinesin function leads to systemic physiological consequences of disrupted intracellular trafficking. Kinesin 1 knockdown significantly impaired larval development from first to third instar, showing greater than 40% reduction in klc animals and 75+% reduction in khc. Previous reports from khc and klc mutant animals describe paralysis and stalled larval development during the second instar stage ^67–69^. Collectively, kinesin 1 knockdown animals are severely impaired developmentally and functionally by the third-instar larvae stage, while minimal impacts were observed for kinesin 3 knockdown. To further dissect the molecular and cellular underpinnings of these effects a thorough assessment of neuromuscular transduction was conducted.

Motor neurons (MNs) projecting from the VNC to body wall muscles not only propagate electrical signals which dictate the patterned output regulating muscle contractile force/timing underlying larval movement, but also transport cargo and numerous other cellular material necessary for neuronal growth, development, and homeostasis ^38,71^. The glutamatergic MNs-Ib (tonic) and MN-Is (phasic) subtypes within the VNC innervate body wall muscles in highly stereotyped morphological varicosities with reproducible branching patterns and terminal-specific bouton numbers along the muscle surface ^49^. Knockdown of khc significantly reduced the extent of innervation along the muscle surface for MN-Ib, and a concomitant reduction in the number of boutons. Interestingly, these effects were not observed for the MN-Is subtype. This difference could arise because of developmental effects, axonal-retraction/degeneration, or a failure of homeostatic plasticity to maintain synaptic structural integrity ^71,72^. Type Ib terminals have greater mitochondrial density and demand 2x the ATP per SV fusion event ^73,74^.

Consequently, the metabolic demands of the Ib-terminals may make them more susceptible to failure in transport of critical cargo like mitochondria ^75^. Noteworthy, our previous work demonstrated that MN-Is terminals precede MN-Ib terminals developmentally which could contribute to the morphological distinction observed herein^71^. A definitive role for kinesins in axonal/growth cones pathfinding and proper development or maintenance of synaptic connections remains unclear ^24,76^. A reduction in innervation length or bouton number for unc^104^ was not observed, however, knockdowns of both kinesin 1 and 3 resulted in a reduction in BRP density at MN terminals (# of brp/bouton). A reduction in innervation length, bouton number, and BRP density correlated with a profound reduction in EJP amplitude, reported previously ^69^. The lack of effect of kinesin 3 knockdown was initially surprising given previous reports in the field where unc^104^ bris reduced EJP frequency ^77^. However, the lack of observable morphological changes in synaptic structure does strongly correlate with a wild-type EJP amplitude. A reduction in BRP density in MN-Is terminals in unc^104^ knockdown animals was observed, but previous reports suggest that this is insufficient to cause quantifiable changes in EJP amplitude as excess BRP exists at AZs ^50,78^.

Changes in EJP amplitude reflect alterations in synchronous NT release from SVs following depolarization ^79–81^, which likely result from the structural changes at the NMJ. Fascinatingly, another striking observation from both kinesin 1 and 3 knockdown animals was the dramatic reduction in NP abundance at the synaptic varicosities along the surface of body-wall muscles. The cumulative impacts of axonal aggregates and trafficking deficits resulted in a 75% reduction in NP abundance, and ∼90% reduction in NP moving through boutons. The effects were even more severe when examining synaptic capture and high-frequency-indued NP mobilization and release, where neither NP transiting from axons into bouton was observed, nor was a significant drop in GFP fluorescence observed following 70 Hz stimulation ^51,82^. Noteworthy, live imaging from boutons revealed no mobility in NP. Such a dramatic impact on NP release has been reported previously in the absence of calcium, however, to our knowledge, no other mechanism has been demonstrated to completely inhibit NP release from boutons ^83^. The lack of DCV mobility and release from boutons may be a consequence of inadequate ATP from insufficient mitochondrial trafficking, a completely incapacitated kinesin motor, or an NP that has uncoupled from the cytoskeleton.

Motivated linear crawling is the most highly stereotyped and orchestrated repetitive patterned behavior, produced from nearly identical contraction waves in body wall muscle of *Drosophila* larvae ^84,85^. Muscles directly below the cuticle contract, then relax, to propagate peristaltic contraction waves from the posterior to the anterior of the larvae, predominately in abdominal segments ^38^. Our lab developed a robust force transducer assay to assess how changes in neuromuscular transduction manifest as alterations in excitation-contraction coupling (ECC) by examining changes in muscle force production with a 10 μN resolution ^38^. Knocking down khc severely impaired larval force-production such that no observable force was generated until 20 Hz stimulation, and ∼80 % reduction in muscle force production was observed. The reduction in muscle force production is likely a product of the impaired neuromuscular transduction we observed in our EJP recordings, as we did not observe any dramatic changes in muscle ultrastructure. This correlates with our previous work that ECC is most strongly coupled to neuromuscular transduction ^38^. Altered synapse morphology, neuromuscular transduction, and ECC in khc knockdowns all correlated with the deficits in larval crawling observed. The reduction in larval crawling velocity, linearity, and linear displacement observed in our locomotory is consistent with previous reports or larval paralysis during the second and third instar stages, as well as *C. elegans* where knockout of Unc-116 (KHC) caused a suppression of overall crawling movement ^86,87^. Knockout mice of KIF5b (Kinesin 1) also show clear impairments in rhythmic locomotory behavior ^88^.

In summary, our data demonstrates which of the kinesin proteins are critical regulators of intracellular trafficking and reveal disruption in expression of kinesin 1 and 3 led to severe downstream implications, even when altered solely in motor neurons. Downstream impacts of kinesin disruption caused significant changes in NP abundance at boutons, and changes in synaptic morphology, including innervation length, bouton number, and active zone composition. Spinning disc confocal microscopy enabled subquantal level resolution of NP at boutons, revealing a profound impact on NP mobilization and release from boutons. Collectively, these disruptions manifest as significant reductions in neuromuscular transduction, excitation-contraction coupling, and larval crawling in kinesin 1 knockdowns, but not for kinesin 3. Additionally, the unique role for kinesin 3 in retrograde signaling may provide evidence for cargo-specific motors to facilitate unique transport requirements. Taken together we have not only identified which kinesins are critically involved in organelle trafficking, but also revealed critical disruptions in cellular morphology, function, physiology, and behavior in genetically disrupted animals.

## Acknowledgements

We would like to thank Professor Troy Littleton (Massachusetts Institute of Technology) for anti GluRIII antibody. Members of the Ormerod lab for helpful commentary on the manuscript. Research supported by the National Institute of General Medical Sciences of the National Institutes of Health under Award Number R15GM155985 to KGO. We also thank generous financial support from Middle Tennessee State University to KGO.

## Supplementary Figures and videos

Supplemental Video 1: Real-time, in vivo intracellular trafficking of fluorescently tagged neuropeptides (OK6-Gal4>UAS-dilp2-GFP).

Supplemental Video 2: Post-processed video using computer software (ilastik) to isolate individual DCVs, then track their position from frame-to-frame.

Supplemental Video 3: Infrared images of OK6-Gal4 third instar larvae captured at 10 frame per second in the absence of ambient light.

Supplemental Video 4: Infrared images of OK6-Gal4<UAS-khc-RNAi third instar larvae captured at 10 frame per second in the absence of ambient light.

Supplemental Video 5: Infrared images of OK6-Gal4>UAS-unc^104^-RNAi third instar larvae captured at 10 frame per second in the absence of ambient light.

